# What’s the Temperature in Tropical Caves?

**DOI:** 10.1101/2020.07.21.213579

**Authors:** Luis Mejía-Ortíz, Mary C Christman, Tanja Pipan, David C Culver

## Abstract

Hourly temperature was measured for approximately one year at 17 stations in three caves in Quintana Roo, Mexico. Thirteen of these stations were in the extensive twilight zones of all three caves. All seventeen stations showed seasonality in temperature with a 3 °C drop during the Nortes season. Two of the caves, Muévelo Sabrosito and Muévelo Rico, showed greater variability during the winter months while in Rio Secreto variability was greatest during the rainy season. Río Secreto is less open to the surface than the other two. All sites also showed a daily temperature cycle, although it was very faint in some Rio Secreto sites. While temperature variability is diminished relative to surface variation, its temporal pattern is worthy of further study.

## Introduction

Besides the absence of light, caves are distinguished from surface habitats by their relative constancy of temperature. For example, Eigenmann [1] divided a cave into three zones—twilight, a region of fluctuating temperature, and the inner cave region of constant temperature, excluding the entrance itself. Gèze [2] proposed that there is a zone of constant temperature *(practiquement invariable)* in most caves. Given this view of invariance of cave temperature, much of the early interest in cave temperature was how it changed from cave to cave with respect to latitude and altitude rather than how it varied within a cave [2,3]. Cave temperature should be the mean annual temperature of the surface [3] although other factors such as air flows can cause deviations. This relative constancy of cave temperature makes the stable isotopic composition of cave speleothems a useful proxy for paleoclimate [4]. Cave temperatures have also been used directly to evaluate shorter term natural and anthropogenically induced climate changes on the scale of decades [5,6].

The relatively low precision of mercury thermometers and their fragility limited the scope of cave temperature studies, a problem which has been solved by the widespread availability of accurate, sturdy digital temperature probes [7,8]. This, together with a growing interest in the physics of cave temperature, has resulted in a large increase in the understanding of our knowledge of cave temperature and its dynamics. Analytical mathematical models appeared early in the 20th century [9] and continue to have a robust presence in the field [10,11,12]. As Cigna [7] points out, a renaissance of cave temperature and climate studies in general would not be possible without an increase in the accuracy and automation of measuring devices, i.e., dataloggers. These physical studies have addressed a series of questions, including:

- Mean temperature prediction using passage size, entrance size, and exterior temperature [13].
- Time lags between exterior and cave temperatures [7,11,14]. These lags are often weeks to months.
- Temperature effects on wall condensation and evaporation [12].
- The relationship between ventilation and temperature [15].

In addition, there are extensive published cave temperature series, some covering multiple years. Most prominent among these are the temperature time series for multiple sites in and near Postojna Planina Cave System (Slovenia) by Šebela and her associates [16,17].

While physicists have been concerned with equilibration of air masses, water, and the surrounding rock [11,12,18] they have not focused on detection of cycles, either daily or yearly, although Stoeva et al. [6] did analyze data for the presence of multi-year cycles correlated with sunspot cycles. With the exception of studies focusing on non-cave shallow subterranean habitats [19], very little attention has been paid to daily, monthly, or seasonal cycles.

We know of no cases where temperatures are truly constant, but the amplitude of variation is reduced at least on an annual scale [20]. In deep cave sites, amplitude of change has been reported to be approximately 1 to 2 °C, as is the case in Karchner Caverns in Arizona [7] and at 1100 m depth in Sistema J2 in Slovenia [11]. Culver and Pipan [19] report on extensive temperature measurements in Pahoa Cave, Hawaii, where annual variation is less than 1 °C less than 100 m inside the cave.

In contrast to physicists, until recently biologists have been largely content to assume temperature constancy, and to make some important assumptions about this constancy. The standard view is that constancy (1) makes cave dwellers highly vulnerable to environmental change because they have had little opportunity to adapt to a varying environment and (2) that neither temperature nor light provide any cues, either daily or seasonally, to set circadian and annual rhythms. The Romanian biologist Emil Racovitza, widely held to have ushered in the modern study of cave biology [21], a very discerning and skeptical chronicler of the science of biospeleology, stated [22,23]:

> *“We can admit that temperature is constant and it corresponds generally to the mean annual temperature of that place in deep caves, in fissured massifs, in phreatic sheets and groundwater.*
>
> *Surely meteorologists armed with ultra-sensitive instruments will discover without a doubt, variation, that in absolute, can be considered important, but relative to the place and their influence on living beings, these variations are lower than those observed in the surface environment, and therefore it is agreed not to bring them into account” …*

After mentioning a few anomalies, such as open air pits, he concludes:

> *“No matter these exceptional facts, we can consider the subterranean environment as a habitat with constant and low temperature, but not identical in all its extent” …*

That is, temperature variation in caves was held to be without biological interest. It is fair to say that this opinion is still widely held by speleobiologists except as it makes the cave fauna vulnerable to climate change [20].

An exception to this lack of interest in temperature variation by biologists is an approach pioneered by Mammola and Isaia [24,25] with their study of niche separation in cave dwelling spiders. In one study, they demonstrated that *Meta menardi* and *Meta bourneti* occurred at different temperatures, but that the differences were sometimes less than 1 °C. Pipan et al. [26] argue that temperature is one of the five key factors that should be measured to assess subterranean ecosystem health and change, in part because it is a surrogate for many more difficult to measure environmental parameters.

### Goals of the Present Study

The advent of dataloggers that measure temperature to the hundredths of a degree and relative humidity to a tenth of a percent allows for a more detailed look at the subterranean environment itself. In spite of the overall reduction in amplitude of variation in subterranean habitats, it is possible to detect differences among closely adjoining sites [16] and to detect the presence of daily and annual cycles [19,26]. In this study, we begin a characterization of the cave environment in terms of temperature. To this end we examine year long temperature records at both photic and aphotic stations, taken hourly, and compare these records, both between different dataloggers in the same cave and among three different caves in Quintana Roo, Mexico. Light intensity provides the backdrop for comparisons.

Our purpose in this paper is two-fold. First, we demonstrate that variation in temperature, although damped relative to surface habitats, shows interesting patterns. These patterns are relevant not only to biologists studying cave life, but to paleoclimate studies [4] and climate change heat transfer in the environment [18]. Second, we suggest some protocols for the analysis from data loggers put in caves and other aphotic habitats.

## Methods and Materials

### The Study Caves

The three caves are located in the Quintana Roo in the Yucatan Peninsula (Fig 1) in an area, with one of the highest densities of cave passages (mostly flooded) in the world [27,28]. Air filled caves are also numerous and they are constrained to a relatively thin layer of flat-bedded limestone with a depth of 5 to 10 m to the water table, and a surface topography of gentle ridges and swales with an overall relief of 1-5 m [28,29]. The area has an annual cycle of precipitation characterized by three seasons: Nortes (cold front season between November and February), dry season (March to May), and rainy season (June to October) which is the hurricane season [30]. During the rainy season 70% of the precipitation occurs. The annual mean temperature is 25.8 °C and the overall precipitation at Playa del Carmen averages 1500 mm over a period of 2005 to 2014 [30].

**Fig 1.**
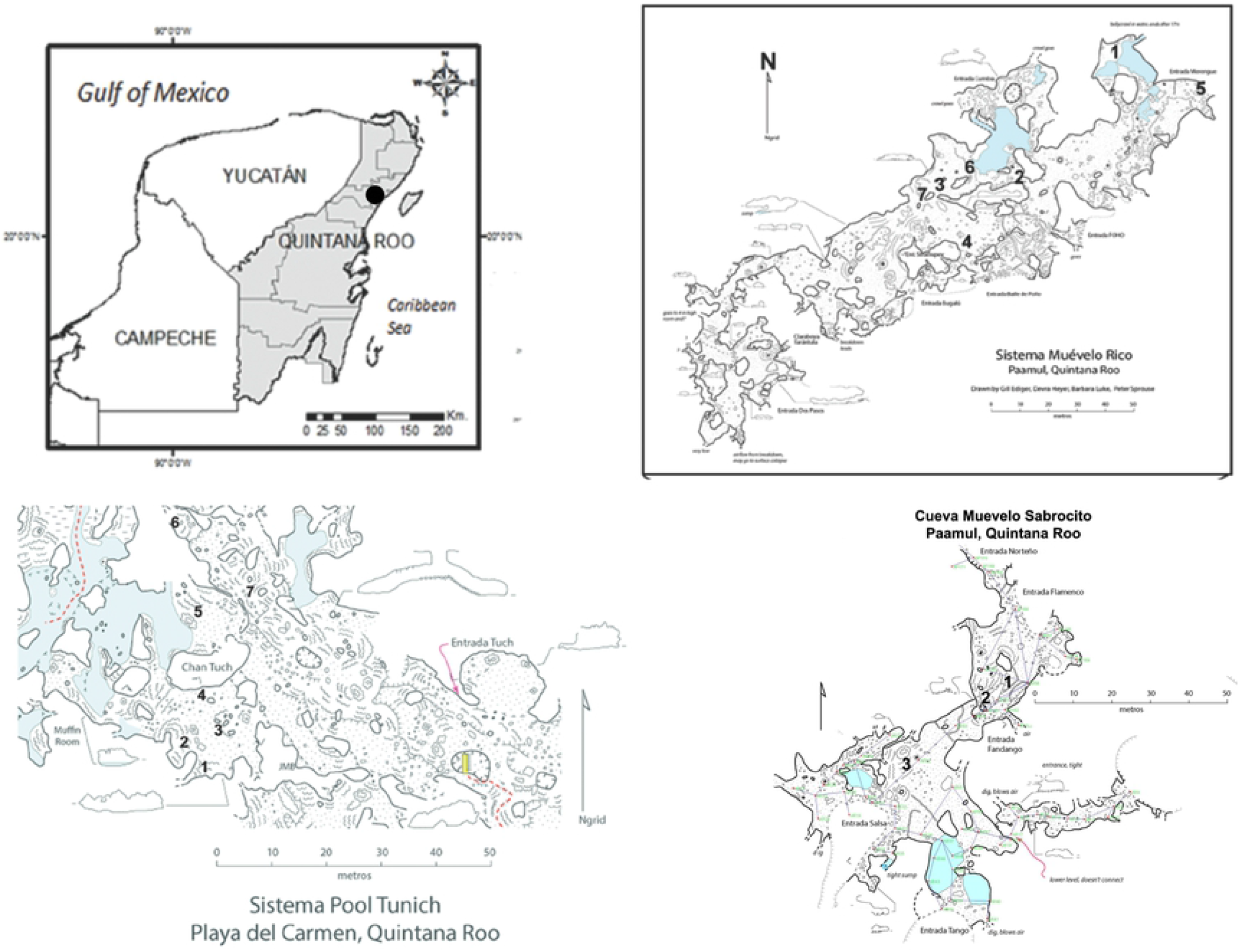
Locator map for caves and sampling sites in the study area. Maps courtesy of Peter Sprouse and colleagues.

Sistema Muévelo Rico (20°32’05.1”N, 87°12’16.5”W) is located near the settlement of Paamul, in the Mexican state of Quintana Roo (Fig 1). Its surveyed length is 1151 m with a vertical extent of only 4 m [27]. Sistema Muévelo Rico has a large number of entrances, more than 12, if skylights are included. Because of the close proximity of the water table to the surface, vertical development and subterranean terrestrial habitats are very restricted. The cave, with an elevation of 7 m at the entrance, is less than 2 km from the Caribbean Sea. It was originally chosen for study by Mejía-Ortíz et al. [31] because of its extensive twilight zone and extremely small aphotic zone. There were seven monitoring points in the cave.

Muévelo Sabrosito (20°53’N, 87°20’W) is a small cave immediately adjacent to Sistema Muévelo Rico. It has six entrances and no aphotic zone. Its surveyed length is 400 m with a depth of 4 m [27]. It has a more open aspect than Sistema Muévelo Rico. There were three monitoring points, all near the Perro Negro section of the cave (Fig 1).

Río Secreto (20°35’27”N, 87°8’3”W) is a shallow, horizontally developed cave with 42 km of surveyed passages. It is a tourist cave and the tours are conducted in a small section of the cave. The main entrance is 5 km from the Caribbean coast and 12 km NE of the other two caves. Tides can affect the water table in Río Secreto up to several cm [30]. There were seven monitoring points clustered in the vicinity of the Tuch entrance (Fig 1).

### Temperature Measurement

Temperature was measured at hourly intervals for the following dates:

- Sistema Muévelo Rico—5 April 2015 to 28 March 2016, n=8593
- Sistema Muévelo Sabrosito—24 September 2018 to 24 October 2019, n=9477
- Río Secreto (Tuch entrance)—25 September 2018 to 26 October 2019, n=9515 Onset Computer Corporation HOBO^TM^ U23 Pro v2 data loggers were used to measure temperature and readings were accurate to ±0.21 °C with a resolution of 0.02 °C.

### Data Analysis

Spectral analysis, periodograms, autocorrelations and partial autocorrelations were done on the hourly data to detect possible cycles. Cycles up to a period period of 600 hours (25 days) were reported. Analyses were done using JMP^®^ Pro 13.2.0 (©2016 SAS Institute, Inc. Cary, NC). Basic statistics (mean and ranges) were done using EXCEL™.

For estimating monthly means, the hourly data were first averaged over each day to obtain daily means for input to the analyses. In addition, for Río Secreto, the sensors were identified as belonging to two sensor groupings depending on whether light was present. General linear models (GLM) with non-constant variance and covariances among observations were used to estimate the monthly temperature means for each cave separately. The model included fixed effects of month within year and sensor group; temporal autocorrelation of the observations was captured by assuming the residuals were correlated according to a autoregressive process with a lag of one (AR(1)); and variance was assumed to differ by month. The AR(1) covariance was chosen because temporal autocorrelations showed a strong value at a 1 day lag but a small partial autocorrelation value for a lag of 2 days (results not shown). It was expected that some months would have more variable values than others. Other fixed effects were also considered, namely season and sensor but were found to be non-informative and statistically non-significant so were dropped from the model.

## Results

### Overall temperature patterns

Basic statistics for the 17 sites in the three caves are shown in Table 1 and Figure 2. Raw temperature data for the three caves are given in S1 Table (Sistema Muévelo Rico), S2 Table (Sistema Muévelo Sabrosito), and S3 Table (Río Secreto).

**Fig 2.**
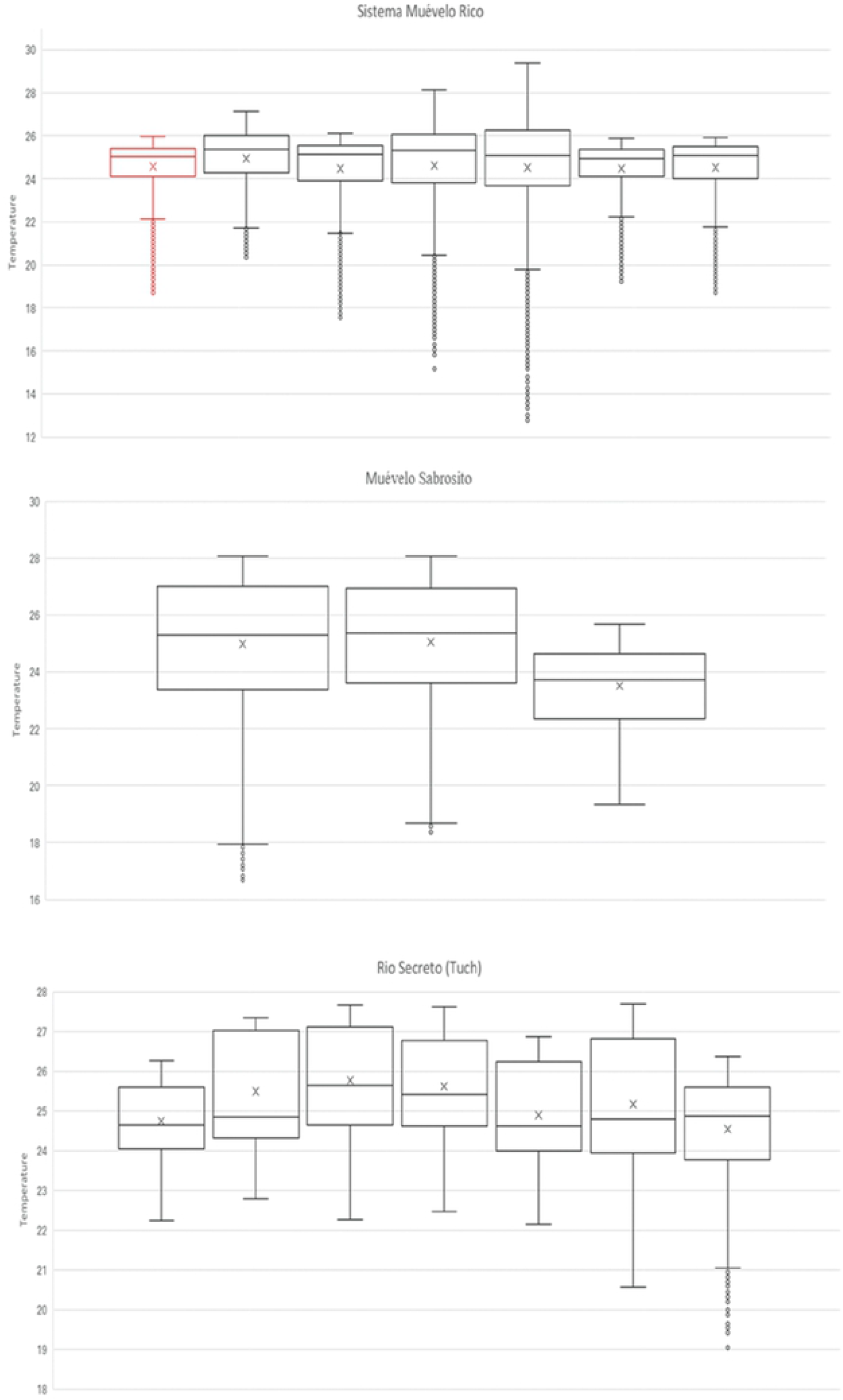
Box and whiskers plots of temperature variation in the individual stations.

**Table 1.** Basic temperature (°C), and light intensities (lux) for the 17 monitoring stations.

| Cave | Station | Temperature |  |  | Light<br>Lux |
| --- | --- | --- | --- | --- | --- |
|  |  | Mean | Min | Max |  |
| Sistema Muévelo Rico | 1 | 24.6 | 18.7 | 30.0 | <0.1 |
|  | 2 | 24.9 | 20.3 | 27.2 | <0.1 |
|  | 3 | 24.5 | 17.5 | 26.1 | <0.1 |
|  | 4 | 24.6 | 15.2 | 28.1 | <0.1 |
|  | 5 | 24.5 | 12.8 | 29.4 | 466 |
|  | 6 | 24.5 | 19.2 | 25.9 | 0.2 |
|  | 7 | 24.5 | 18.7 | 26.0 | <0.1 |
| Sistema Muévelo Sabrosito | 1 | 25.0 | 16.7 | 28.0 | <0.1 |
|  | 2 | 24.7 | 18.4 | 28.1 | <0.1 |
|  | 3<sup>a</sup> | 23.5 | 19.3 | 25.7 | 0.3 |
| Río Secreto (Tuch) | 1 | 24.8 | 22.2 | 26.3 | 0 |
|  | 2 | 25.5 | 22.8 | 27.3 | 0 |
|  | 3 | 25.8 | 22.3 | 27.7 | 0 |
|  | 4 | 25.6 | 25.4 | 27.4 | 0 |
|  | 5 | 24.9 | 22.1 | 26.9 | <0.1 |
|  | 6 | 25.2 | 20.6 | 27.7 | 0.8 |
|  | 7 | 24.6 | 19.0 | 26.4 | 7.7 |

Station 5 in Sistema Muévelo Rico is at the entrance.
<sup>a</sup> sensor moved on 27 January 2018, n=2999

Temperatures at the seven cave stations in Sistema Muévelo Rico had an amplitude of between 6.7 and 12.9 °C, and the three cave stations in Muévelo Sabrosito had a temperature range of between 6.4 and 11.3 °C. In the Tuch section of Río Secreto, where there was a large dark zone (Table 1), temperature ranges at the seven stations varied between 2.0 and 7.4 °C. For all stations in all caves, temperature extremes occurred at the low end, with no high temperature extremes (Fig 2).

Order is by number of the station. X’s indicate means; the horizontal line within the box, the median; the boxes, the inter-quartile range; whiskers, 1.5 times the interquartile range beyond the box; and small circles are outliers.

### Spatial-temporal variation

Plots of temperature through the year are shown in Figures 3 and 4. For all stations, there was a drop in temperature during the winter months of about 3 °C from the summer high. The other seasonal difference was that there was increased variability, usually short term fluctuations, including daily fluctuations, depending on the season. Short term variability was highest during the Nortes season, when a series of cold fronts come through the region [30].

**Fig 3.**
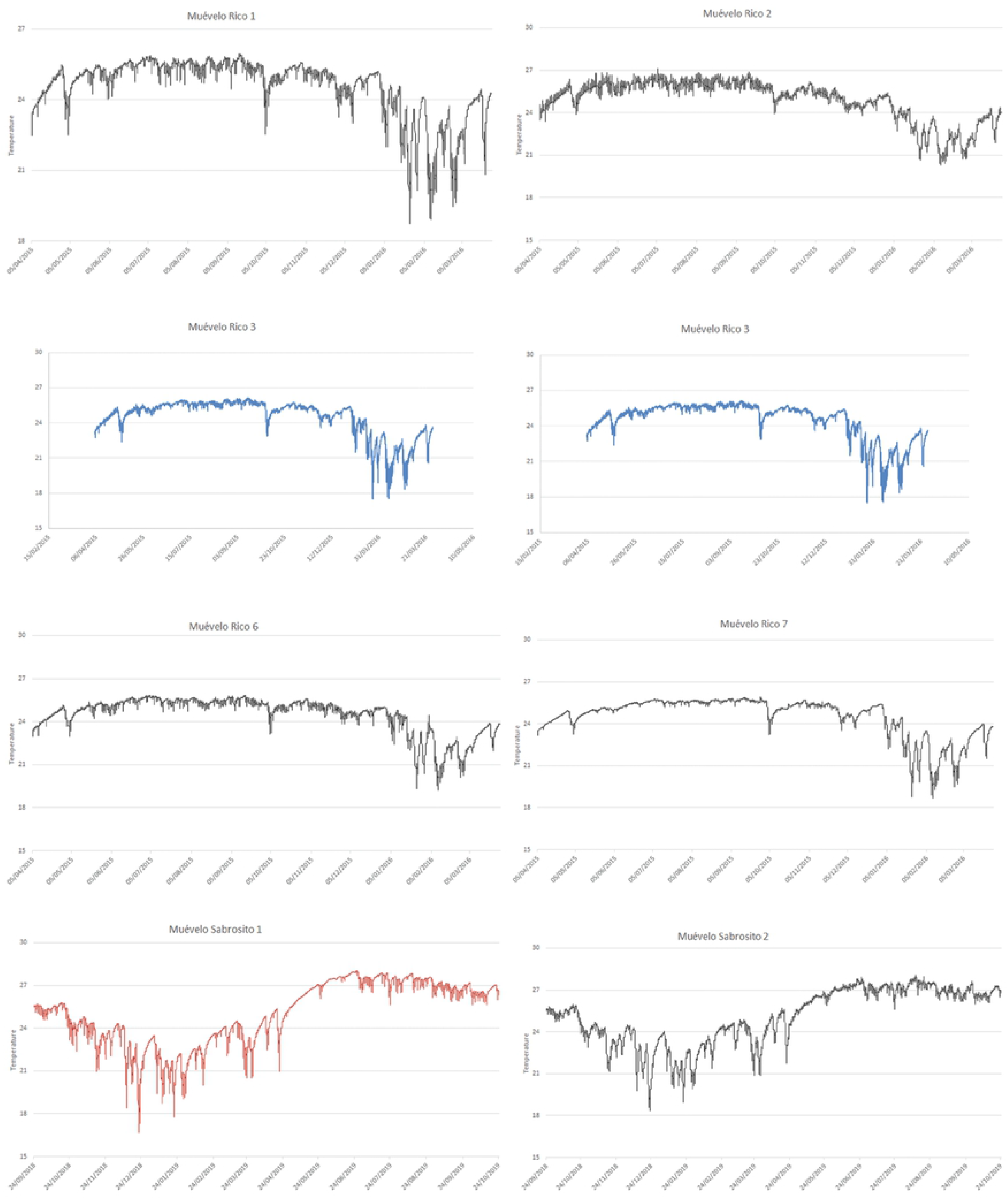
Plot of hourly temperature at stations in Sistema Muévelo Rico and Muévelo Sabrosito. Note that the minimums and higher variabilty all occur in winter.

**Fig 4.**
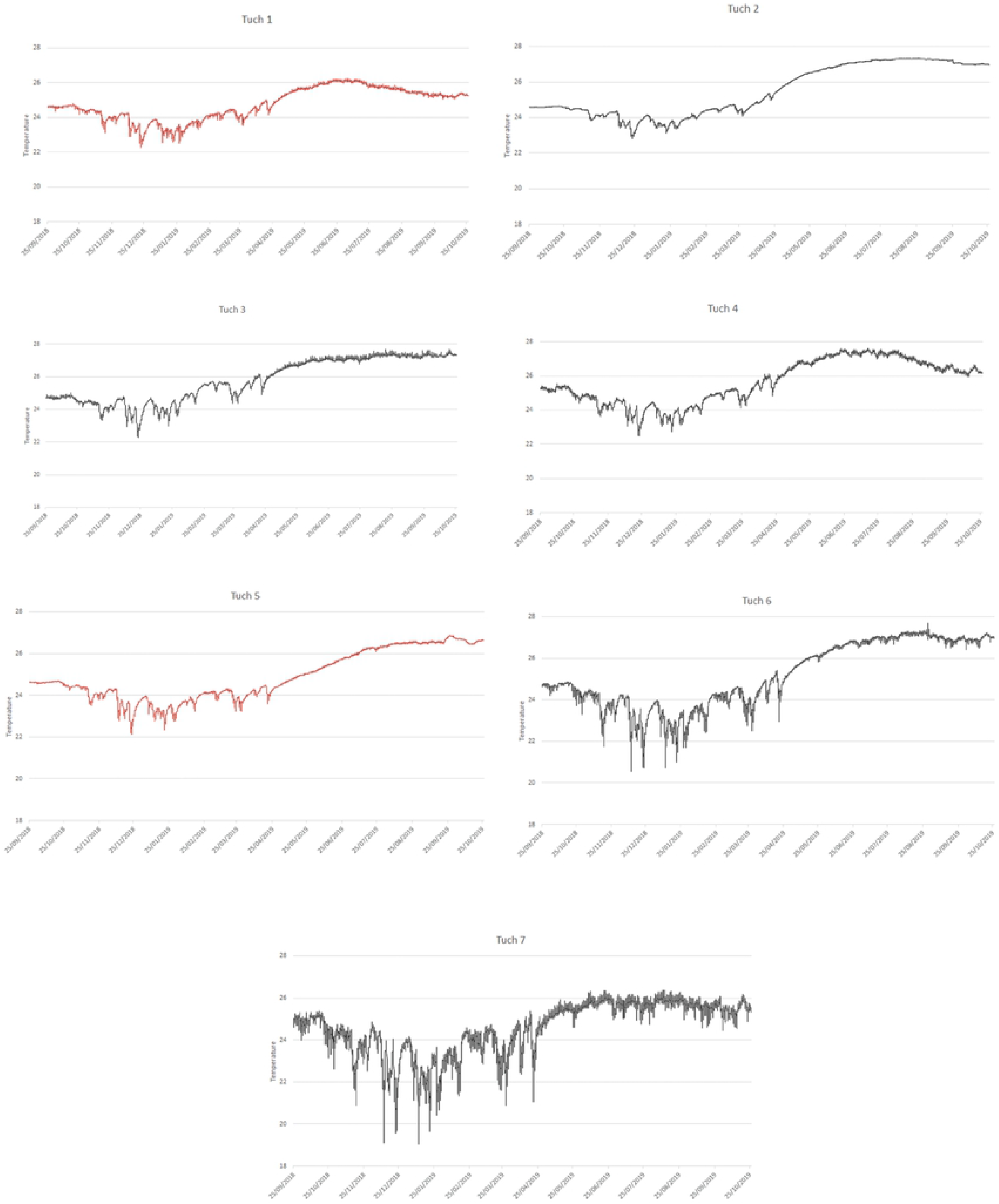
Plot of hourly temperature at stations in Río Secreto near the Tuch entrance. Note that the minimums and higher variability all occur in winter.

For each of the three caves, the best temperature model included month/year fixed effects with allowance for monthly unequal variances, and including autocorrelation with a lag of 1 day. Station (and season) had no effect, which was surprising, especially for Río Secreto, where a dark zone was present. The month/year effect was strong, as can be seen in Figure 5, and was highly significant (p<.0001) in all cases. In addition, the residuals were well-behaved, being approximately normally distributed. For Muévelo Sabrosito and Muévelo Rico, standard errors were less during the summer months.

**Fig 5.**
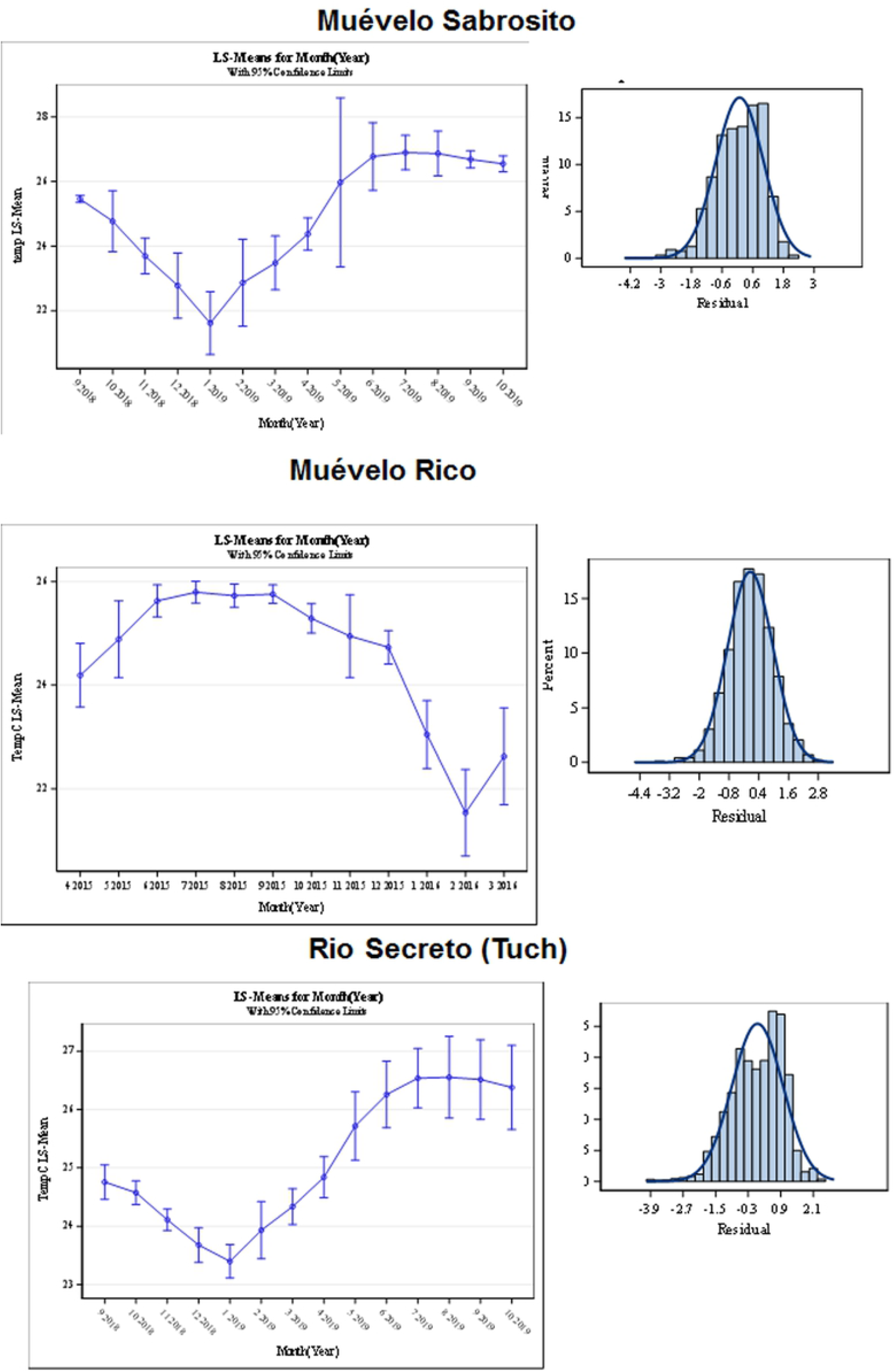
Least Square Means for temperature estimated in the GLM. Left panels are estimates of mean temperature with 95% confidence intervals. Note that the graph for Sistema Muévelo Rico begins at a different month. The entrance station for Sistema Muévelo Rico is not included. Right hand panels are plots of the residuals, compared to a normal distribution.

Since the temperature measurements for Río Secreto and Muévelo Sabrosito were done at the same time, their patterns can be compared. In this case, the interaction of cave x month x sampling site within a cave had a significant effect. When the sites are compared (light and dark sites were grouped separately), the seasonal effects remain the most obvious (Fig. 6).

**Fig 6.**
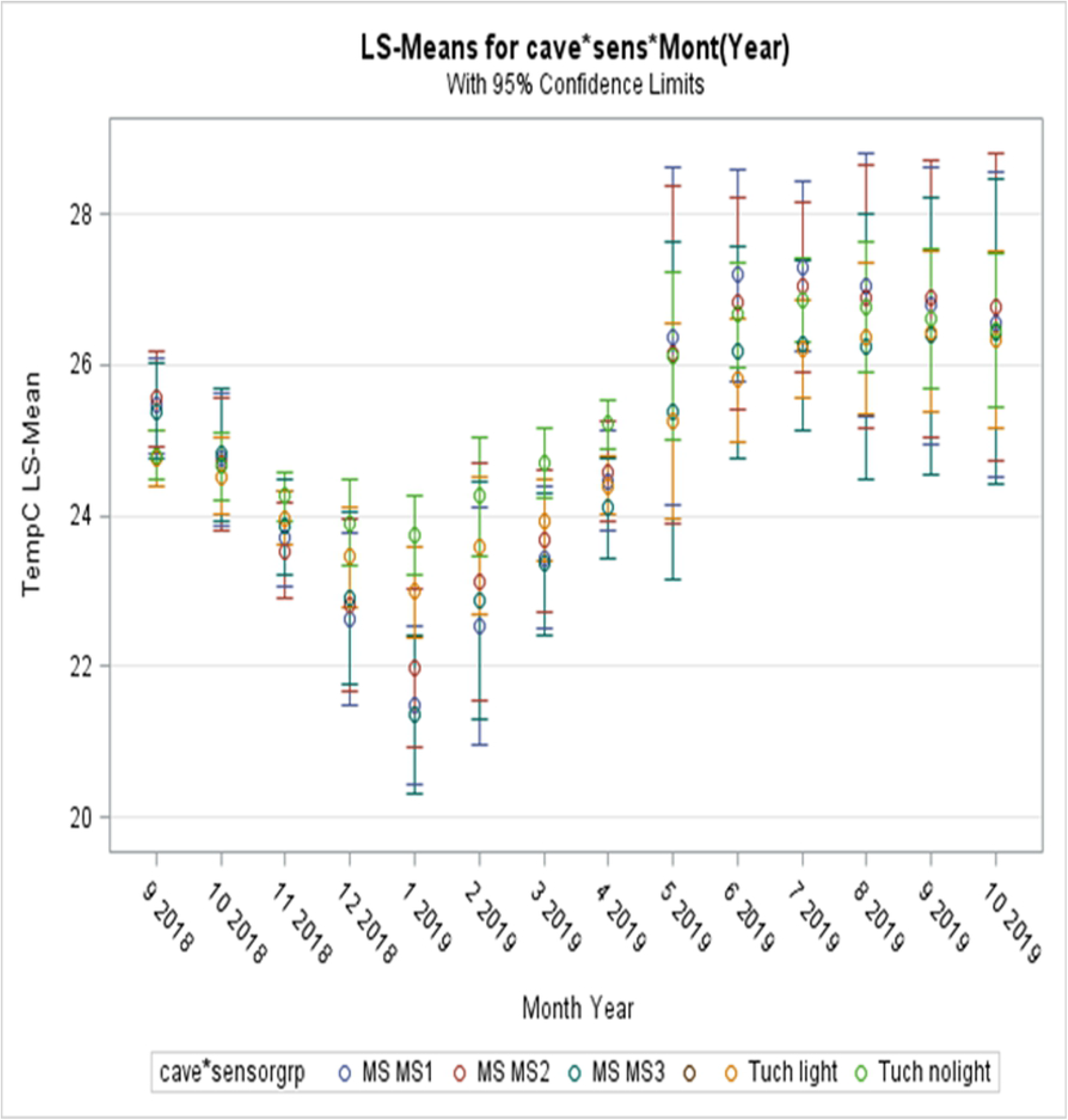
Comparison of Río Secreto (labelled as Tuch in figure) and Muévelo Sabrosito (labelled as MS in figure) based on GLIMMIX model.

One especially interesting comparison is between the dark and light sites in Río Secreto (Fig 7). Dark zone temperatures tended to be higher throughout the year. This relationship would reverse during hotter than normal years.

**Fig 7.**
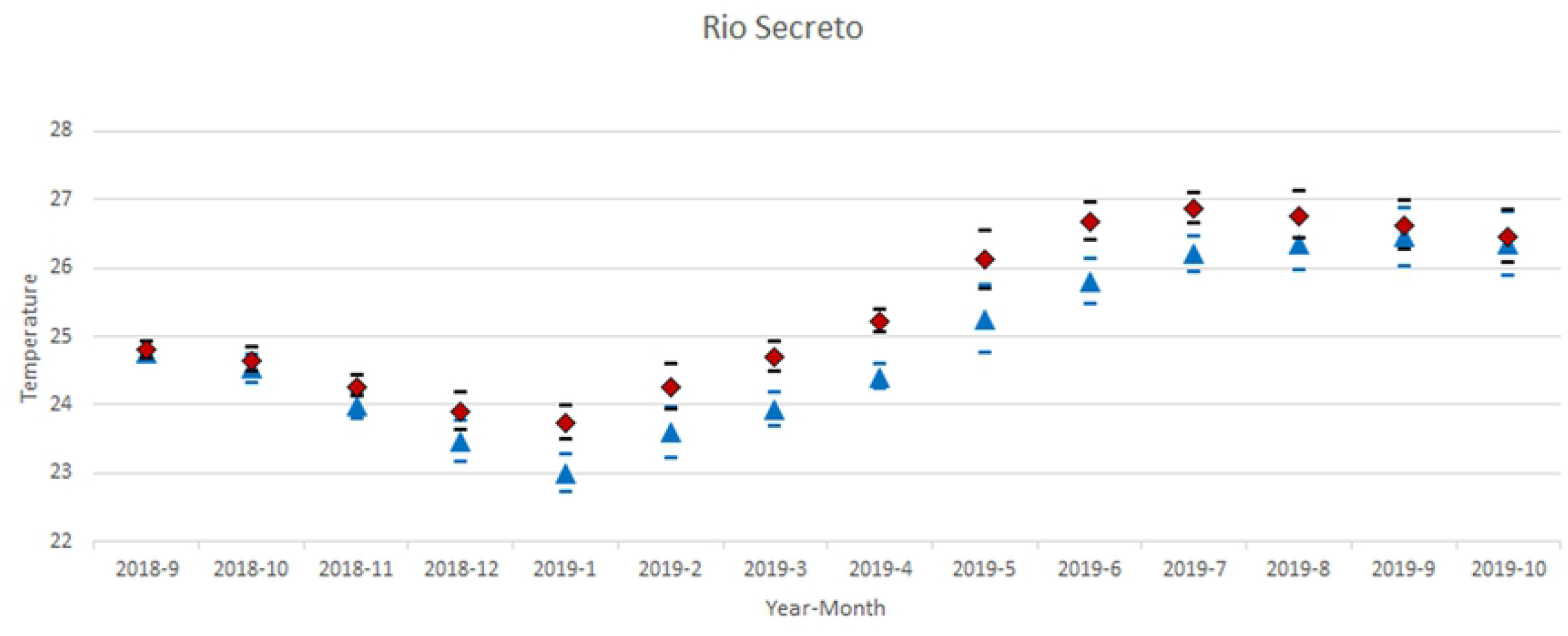
Comparison of GLIMMIX models of aphotic and photic zones in Río Secreto. The diamonds and black error bars are for the dark zone and the blue triangles and error bars are for the light zone. The seasonal cycle is apparent as is the greater variation during the winter months.

### Daily cycles

Spectral analyses of the 17 sites in the three caves are shown in Figures 8–10, for cycles up to 600 hours in length. For all sites in all caves, the pattern was significantly different than white noise (Fisher’s kappa test and Barlett’s K-S test).

**Fig 8.**
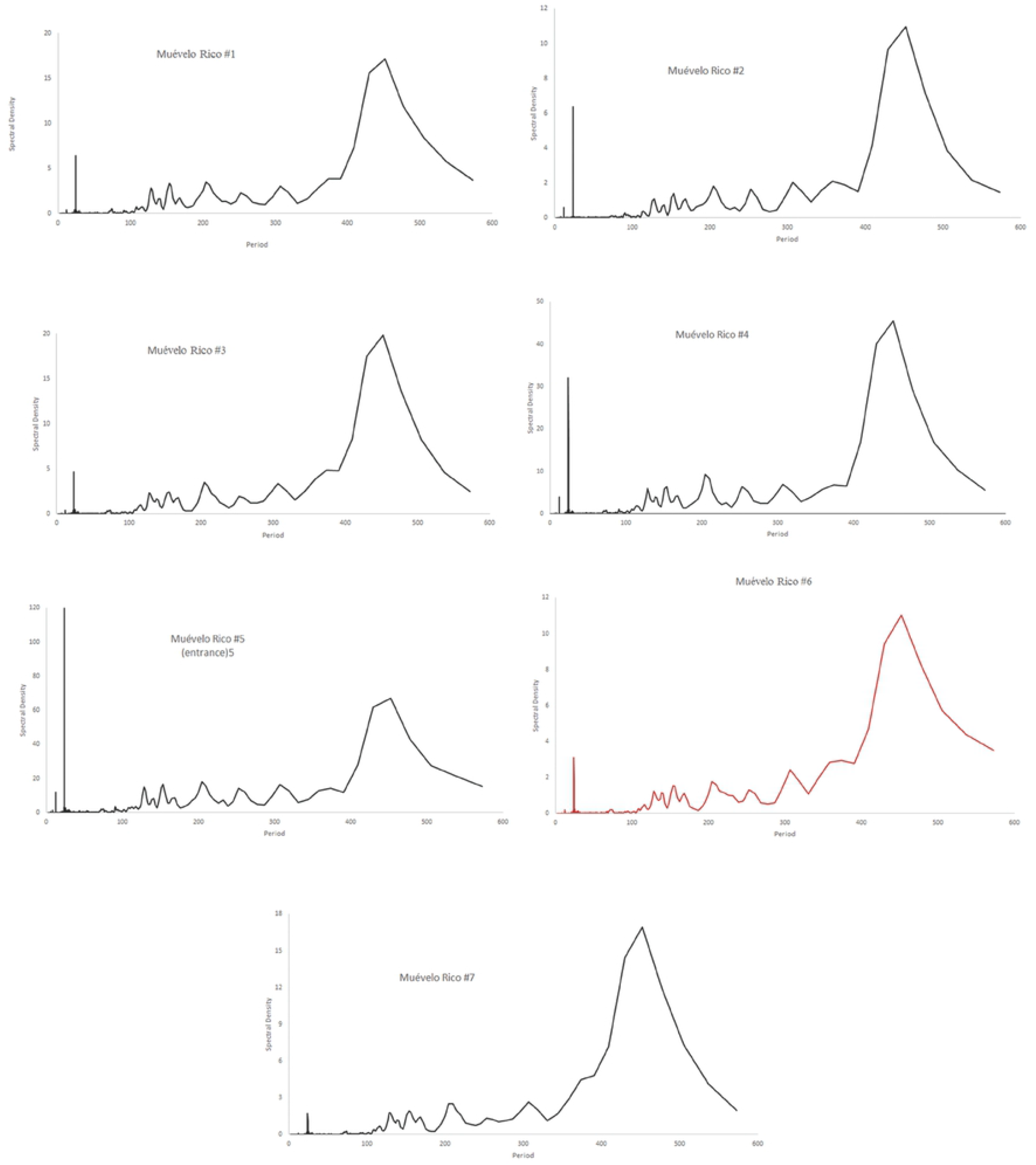
Spectral densities of temperature at the Muévelo Rico sampling stations. Note that the scale of the y-axis is different for different stations.

Sistema Muévelo Rico (Fig 8) and Muévelo Sabrosito (Fig 9) showed a 24 hour spike at all stations, with an odd, much smaller 12 hour spike at several stations (stations 2, 4, and 5 in Sistema Muévelo Rico and station 3 in Muévelo Sabrosito).

**Fig 9.**
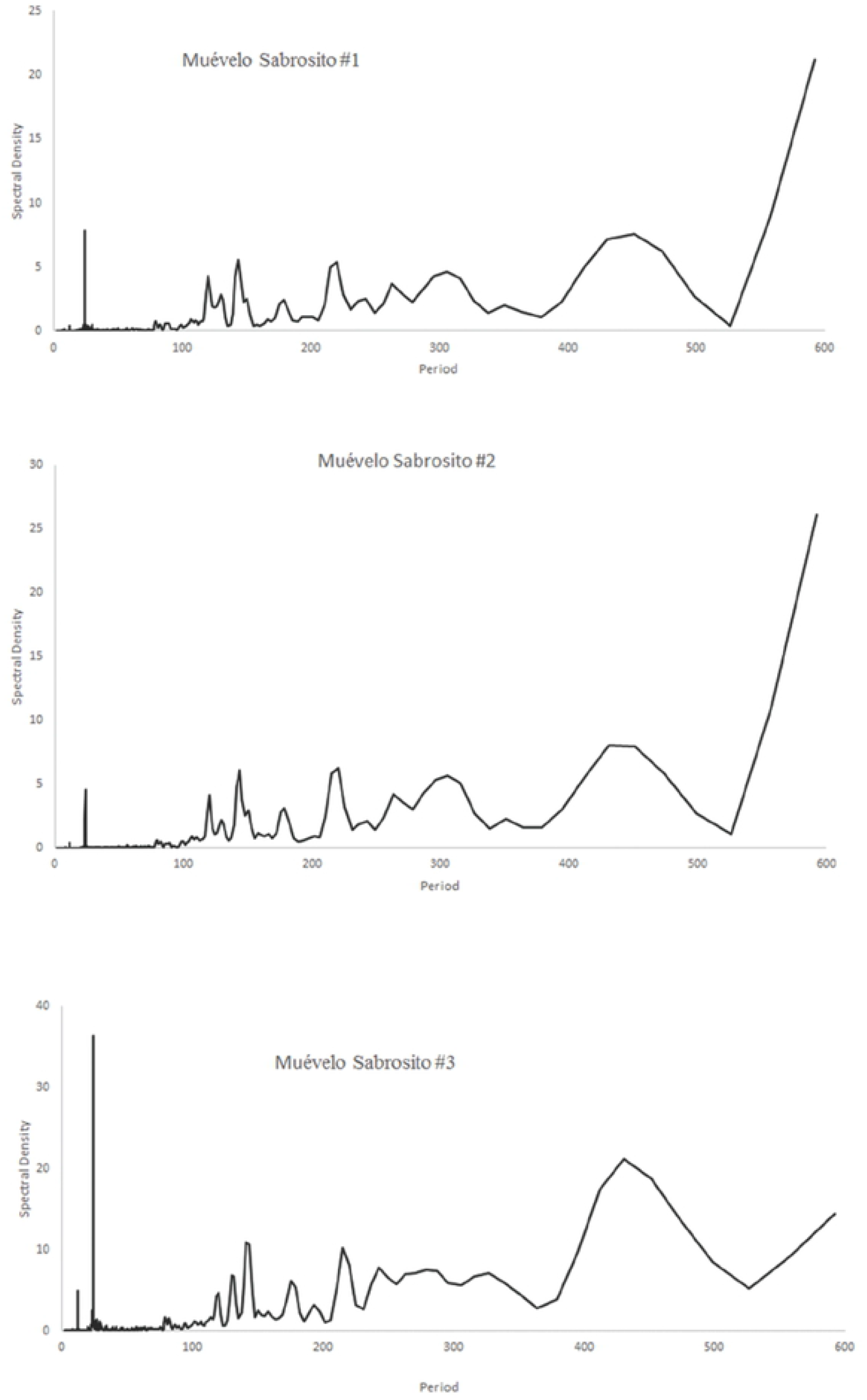
Spectral densities of temperature at the Muévelo Sabrosito sampling stations. Note that the scale of the y-axis is different for different stations.

There was a strong jump in the spectral density at around 425 hours (17-18 days). It was more prominent than the 24 hour spike in all stations except the entrance in Muévelo Rico. In Muévelo Sabrosito, it was equally prominent in #1 and #2, but the 24 hour spike was more prominent in #3.

Río Secreto (Tuch) showed a different pattern (Fig 10). All stations had a 24 hr spike but it was generally weak. The spike in Tuch #2 was barely evident. The 425 hour spike was also less prominent and not really clear in Tuch #3 and #5 where it became bimodal. There was a more or equally prominent jump at 300 hours (12-13 days). Note these were run at the same time as Muévelo Sabrosito.

**Fig 10.**
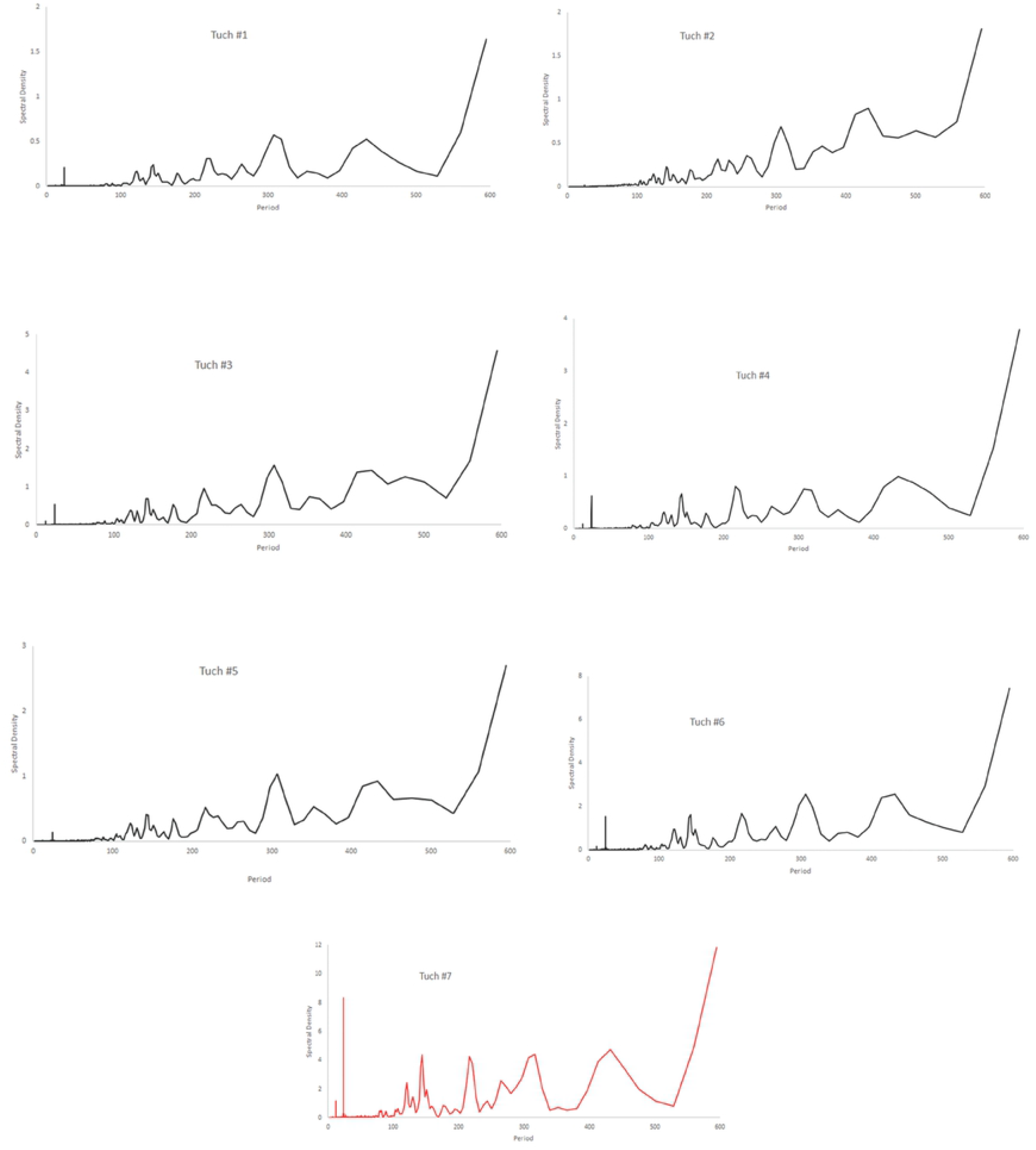
Spectral densities of temperature at the Río Secreto (Tuch) sampling stations. Note that the scale of the y-axis is different for different stations.

## Discussion

### Temporal patterns

In all three caves there was a marked seasonal effect in temperature, and all stations showed a daily cycle of temperature although the signal was extremely faint in stations 2 and 5 of Río Secreto (Fig 10). None of the stations in Río Secreto were particularly deep in the cave, and it seems certain that deeper sites would have no discernible daily temperature cycle. However, it also seems unlikely that there is anywhere in this river cave without temperature variation throughout the year. For example, there is a detectable annual temperature cycle in Karchner Caverns, Arizona, with an amplitude of less than 2 °C [4,7]. Fairchild and Baker [4] outline the many factors that result in spatial and temporal differences in temperature in caves. This variation is of considerable importance to climatologists attempting to use speleothems as proxies for climate change. The rate of equilibrium of cave temperature has been measured at 0.04 °C per year in Villars Cave in France [32], indicating equilibration takes a rather long time. Time lags also imply a different phase for surface and cave temperature cycles [16].

We would expect different caves in different regions to have different patterns with respect to the temporal variation and co-variation of light and temperature. The caves we studied were very shallow with numerous surface connections making variation in temperature on the surface a strong signal in the caves. We do not have detailed surface temperature data available. The estimated monthly means for the interior of Río Secreto had a range of 21.5 to 25.8 °C. By contrast, surface monthly temperatures in nearby Playa del Carmen ranged from 21 to 30°C for a 14 year period [30]. In this case the percent reduction in amplitude was 52 percent. The surface and cave data sets are not strictly comparable but do give a sense of the amount of attenuation of temperature. It is likely that in temperate regions, with greater surface variation, that the attenuation in caves is greater.

### Is the temperature variation observed biologically significant?

We also do not know what the biological response, if any, is to the relatively small amplitude of temperature variation. Large-scale differences in temperatures may well be lethal to many cave organisms since they do not encounter large scale changes. Smaller differences may be important in niche separation, as Mammola and Isaia [25,26] show for subterranean spiders.

There is also a disconnect between lux values observed in caves and shallow subsurface habitats with the lux values typically used in experiments with subterranean animals [33]. The scarcity of surface dwellers and the relative abundance of cave animals in dimly lit Muévelo Rico [31] suggests that very low light levels are in some ways equivalent to no light, at least in terms of faunal composition. Low temperature variation may likewise be equivalent to no temperature variation for the cave inhabitants.

It is well documented that for species limited to caves and other aphotic habitats, that eyes and pigment tend to be reduced or absent compared to related surface-dwelling species [34]. A similar reduction with respect to thermal tolerance may be expected for species living in environments that are nearly thermally constant, such as caves [20]. One hypothesis is that the thermal tolerance of cave-limited species should correspond to the temperature variation in the subterranean habitats where the species are found. The actual pattern, and it is hard to even find a pattern, is quite different [20]. There are a very few cases where temperatures out of the range of those encountered have been reported to be lethal, most notably two species of *Proasellus* isopods living in caves and springs in the French Jura Mountains [35]. However, a third species showed broad thermal tolerances. A more common finding is that thermal tolerances of subterranean species are less than species found on the surface but that the range of thermal tolerance is much greater than the temperature range currently encountered by the species [36,37,38]. Nearly all the studied examples are from temperate zone caves and the situation may be different in tropical caves, where surface variation intemperature is less, where phenotypic plasticity may be reduced in general [39]. The effect of reduced thermal variation may well have other effects, such as a reduction in phenotypic plasticity, but this has been little studied.

The possible reasons for this confusion of results are many. First, testing conditions vary, and it may well be that there are long term effects of temperature change not detected in the experimental protocol. Second, genes for thermal tolerance may have pleiotropic effects, a common situation for eye and pigment loss [40,41]. This is likely the case for heat shock proteins [42]. Third, there may not have been sufficient evolutionary time for the thermal tolerance to attenuate.

### The utility of temperature measurements

The widespread availability of dataloggers to measure temperature provide an unprecedented opportunity to understand the physical environment of caves. At present there is a gap between the views of biologists and physical scientists studying caves— biologists stress constancy and physical scientists stress differences. What is clear from the analyses presented here is that there is a muted variability in caves, but variability with a rich temporal and spatial pattern, even in supposedly constant tropical caves. The mapping of organisms onto this pattern should provide new insights into the ecology of cave organisms.

It is also clear that a simple description of variability does not capture the patterns, especially of temperature. It is not the total amount of variability that is likely to be important, but rather its temporal pattern. We found generalized linear models especially useful for the analysis of this pattern and there are of course other statistical tools available. The point is not to rely on overall measures such as means and standard deviations.

## Acknowledgements

Benjamin Schwartz, Christian Martinez, Ximena Rosales, Jesus Cupul, Alex Contreras, VanessaTafoya, Rodrigo Cisneros, Fernanda Lases, and Raúl Padilla assisted with field work. Alberto Rivero gave permission to visit the Sistema Muévelo Rico and Muévelo Sabrosito. Tania Ramirez gave permission to visit the Tuch entrance of Río Secreto.

## Supporting information

**S1 Table. Hourly temperature data for Sistema Muévelo Rico.**

**S2 Table. Hourly temperature data for Muévelo Sabrosito.**

**S3 Table. Hourly temperature data for Río Secreto (Tuch entrance).**

